# Climate drives geographic variation in individual *Peromyscus leucopus* immunity against zoonotic disease

**DOI:** 10.1101/2024.08.23.609392

**Authors:** Vania R. Assis, Kailey McCain, Rachel A. Munds, Allison M. Brehm, John L. Orrock, Lynn B. Martin

## Abstract

Geographic variation in host immunity could have major influences on disease dynamics, including zoonotic forms that affect humans. Such variation in immunity could be driven by variation in climate, either directly or, more likely, indirectly via resource availability. We compared the immune gene expression of wild *Peromyscus leucopus* mice, the primary reservoir for the bacterium that causes Lyme disease, *Borrelia burgdorferi*, among eight sites spanning 1,400 km of the northeastern United States. We discovered that climate conditions at sites strongly predicted immunity to the most common zoonotic pathogen in the U.S.: mice from warmer, wetter sites were more prepared to resist *B. burgdorferi* infections. Our results reveal a novel pathway by which climate change could affect pathogen spillover or zoonotic epidemics generally.

## Introduction

Zoonoses are pathogens that can be transmitted among wild animals and humans (Chomel 2014). Whereas we have started to gain evidence that host populations can vary immunologically across their geographic range (Bonneaud et al. 2012; Becker et al. 2020), it is yet obscure how this variation is related to particular zoonotic infections. Interactions among hosts, vectors, and pathogens are complex, but host competence, the ability of an individual host to transmit a pathogen to another host or vector (Ashley and Demas 2017; White et al. 2018; Martin et al. 2019), probably plays a major role in cycles and outbreaks of zoonoses (Ostfeld et al. 2014; Albery and Becker 2021). Traditionally, variation in competence has been expected to be higher among species than within species. Growing evidence, though, exists to the contrary (Barron et al. 2015; Martin et al. 2019), and much intraspecific variation in immunity appears to be driven by environmental factors (Gluckman et al. 2005; Martin 2009). For instance, local climatic conditions (Orrock and Danielson 2009) can affect host immunity by modifying resource quality or quantity (Ostfeld et al. 2006; Dhawan et al. 2018). Host individuals might be more prone to exposing themselves to vectors or pathogens as they seek sufficient resources to build and maintain sufficient defenses (Beldomenico and Begon 2010). Natural and anthropogenic stressors might also alter aspects of immunity (Martin 2009; Abolins et al. 2018), potentially changing the roles that individual hosts play in local disease dynamics (Murone et al. 2016; Gervasi et al. 2017). Climate, too, can affect host immunity, particularly in ectotherm hosts and their directly transmitted pathogens (e.g., amphibians and their fungal infections) (Cohen et al. 2020) and vectors that transmit malaria-causing parasites and a variety of viruses (Caldwell et al. 2021). Surprisingly, little is known about climate effects on immunity in endothermic hosts, but small endothermic hosts might well be affected immunologically by climate due to their high metabolic demands and comparative lack of thermal inertia.

To determine whether host populations can vary in their disposition to exacerbate zoonotic risk (i.e., act as superspreaders) (Paull et al. 2012; Martin et al. 2019; Irving et al. 2021) and determine whether such variation is related to local conditions (Altizer et al. 2018; Bouchard et al. 2019; Halliday et al. 2023), we focused on climate effects on hosts of the most common zoonoses in the United States, Lyme disease (Mead 2015; Zinck et al. 2023). Lyme disease is caused by the spirochete bacterium *Borrelia burgdorferi*, transmitted through the bite of blacklegged ticks (*Ixodes scapularis*) (de la Fuente et al. 2017; Kurokawa et al. 2020), and causes over 450,000 human cases each year (Centers for Disease Control and Prevention (CDC) 2024). Blacklegged ticks become infected with *B. burgdorferi* via blood meals from vertebrate hosts, especially certain small mammals (Kurokawa et al. 2020). The white-footed mouse, *Peromyscus leucopus*, is probably the most important reservoir of Lyme-causing bacteria in northeastern North America, partly because this host species can harbor *B. burgdorferi* for long periods without appreciable effects on its health and fitness (Ostfeld and Keesing 2000; Barbour 2017; Milovic et al. 2024).

Here, we studied immune gene expression in *P. leucopus* at eight sites (Fig.1A) of the National Ecological Observatory Network (NEON). These sites spanned the upper Midwestern and northeastern U.S., capturing significant variation in climate (Dantzer et al. 2023) but also occurring where human Lyme disease cases, *P. leucopus*, and the prevalence of *B. burgdorferi* in ticks are high. We used small tissue samples from wild-caught mice to determine whether immune gene expression varied among wild-caught *P. leucopus* and was related to climate at sites (Paull et al. 2012; Estrada-Peña et al. 2014). We focused on variation in immune gene expression in pinnae (ear tissue) samples, in particular, as ticks primarily bite and thus transmit *B. burgdorferi* to this body area (Ostfeld et al. 1993; Råberg 2012; Zinck et al. 2023). We also chose immune gene targets based on their roles in *B. burgdorferi* resistance and tolerance. We chose genes related to resistance and tolerance because these responses to infection have very different ecological consequences (Martin et al. 2016; Adelman and Hawley 2017). Whereas highly resistant animals should reduce the overall risk of transmission of *B. burgdorferi* from mice to ticks, tolerance responses could potentiate transmission by increasing the time a vertebrate host has to transmit *B. burgdorferi* to a tick and/or the number of total ticks that could become infected (Råberg 2014; Martin et al. 2019). Immunologically, resistance to *B. burgdorferi* involves discrimination of the pathogen from host cells via Toll-like receptor 2 (TLR2) (Tschirren et al. 2013). Once bound by *B. burgdorferi* lipoproteins, TLR2 triggers an immunological cascade, including the cytokines interferon (IFN)-γ and interleukins (IL)-1 and 6 (Strle et al. 2011). These cytokines stimulate local leukocytes to clear or at least reduce bacterial burden. Coincidently to resistance gene expression, hosts can initiate bacterial tolerance. For *B. burgdorferi*, this endpoint is partly accomplished by the anti-inflammatory cytokines, IL-10, tumor growth factor-β (TGF-β), and gata-binding protein 3 (GATA3) (Jackson et al. 2014; Martins et al. 2019).

**Figure 1.**
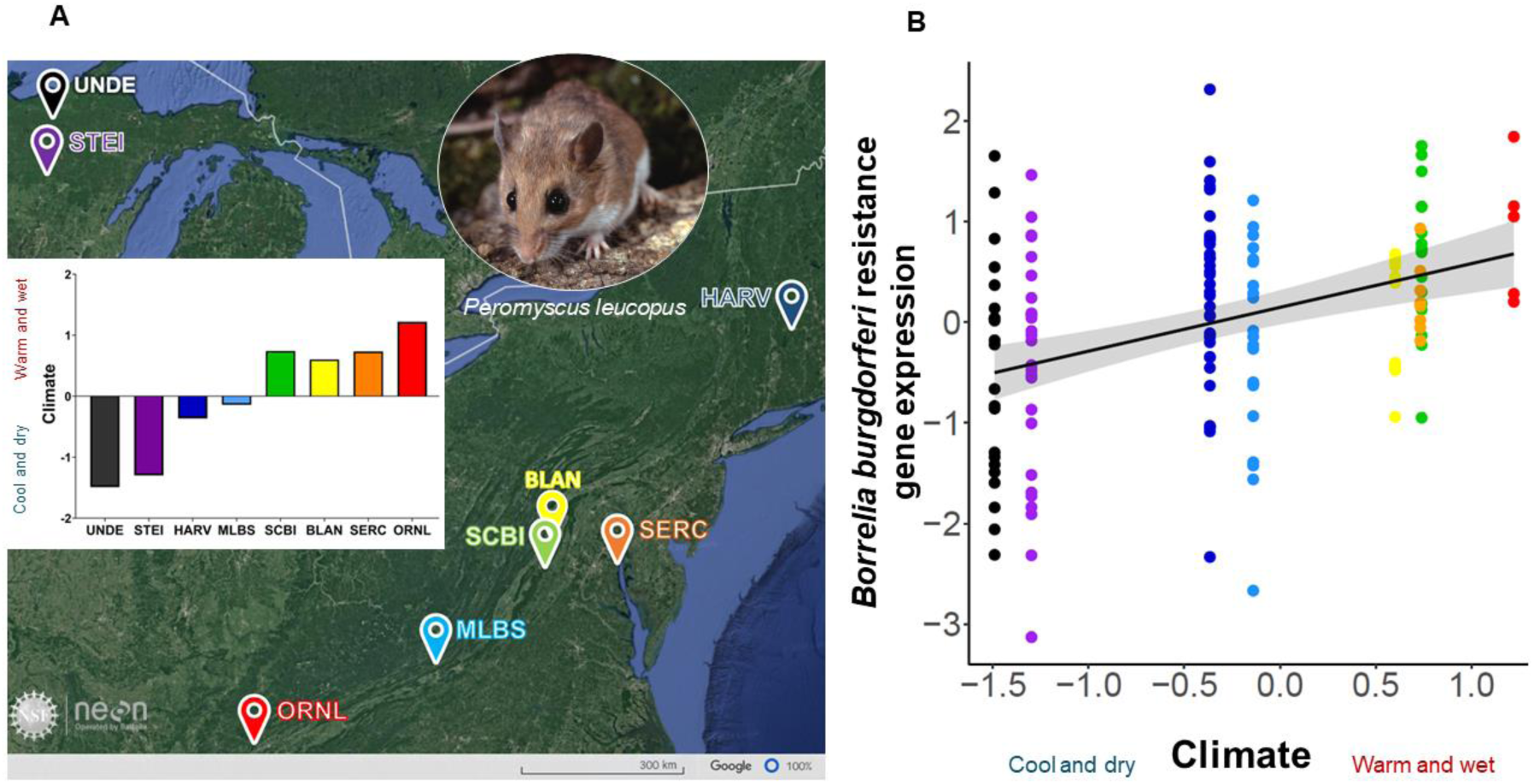
*Peromyscus leucopus* from warm, wet NEON sites have immune systems more poised to resist *Borrelia burgdorferi* infection than mice from cool and dry sites. (**A**). Location of NEON sites involved in this study, all of which have comparatively high incidence of Lyme disease and large *Peromyscus leucopus* populations. Inset figure depicts climate variation among sites with values on the y-axis reflecting scores from a principal component analysis describing annual mean temperature and relative humidity, total annual precipitation, and monthly variation in temperature, relative humidity, and precipitation (unpublished data). (**B**) *Peromyscus leucopus* mice in warm and wet environments show higher resistance immune gene expression than those in cool and dry areas. Values on the y-axis reflect scores from an immune principal component analysis derived from TLR2, IFN-γ, IL-6, and IL-10 gene expression data. Values on the x-axis reflect climate variation scores (unpublished data). Black line represents mean values, and the gray shaded area represents 95% confidence intervals. Abbreviations are as follows: **UNDE:** University of Notre Dame Environmental Research Center, Michigan. **STEI:** Steigerwalt-Chequamegon, Wisconsin. **HARV:** Harvard Experimental Forest & Quabbin Watershed, Massachusetts. **MLBS:** Mountain Lake Biological Station, Virginia. **SCBI:** Smithsonian Conservation Biology Institute, Virginia. **BLAN:** Blandy Experimental Farm, Virginia. **SERC:** Smithsonian Environmental Research Center, Maryland. **ORNL:** Oak Ridge National Laboratory, Tennessee. Map generated on Google Earth. Mice image modified from wildlife photographer and writer, Dr. William J. Weber (available through iStock).

We hypothesized that mice from sites with high mean temperature and precipitation would exhibit the most resistant immune gene profiles (i.e., high TLR2, IFN-γ, and IL-6 expression), either because mice from these warm, wet areas would be least food-restricted and hence capable of erecting a robust immune response to *B. burgdorferi* and/or because tick prevalence (and thus *B. burgdorferi* exposure risk) would be greatest at these sites, evolutionarily selecting for high, constitutive expression of these genes. In contrast, at colder, drier sites, where tick exposure risk and/or food availability would both be lower, we predicted mice would likely have more tolerant immune profiles. Our results generally support these predictions but reveal that climate effects are strong for *B. burgdorferi* resistance but absent for tolerance. Climate might, therefore, underpin Lyme disease hotspots by altering the ability of individual mice to resist infection.

## Materials and Methods

### Animals and Study Site

Pinnae (ear tissue) samples from *Peromyscus leucopus* (*N* = 148) were collected by NEON staff in 2022 from eight sites (Fig. 1A). Immediately upon collection, ear samples were placed in labeled 2mL cryovials and kept on dry ice in the field to prevent RNA degradation. Subsequently, these samples were stored at −80°C at NEON facilities. In November 2022, all samples were shipped to the University of South Florida (USF) overnight on dry ice. The samples were maintained at −80°C upon arrival at USF until RNA extraction. We measured gene expression in ear biopsies partly because ticks are usually attached to pinnae (Ostfeld et al. 1993; Fellin and Schulte-Hostedde 2022), but also because previous studies found a positive relationship between *Borrelia burgdorferi* load in ear tissue and transmission success from host to feeding ticks (Råberg 2012; Zinck et al. 2023) and because this protocol is part of routine sampling by NEON staff. We selected the eight NEON sites specifically because Lyme disease is prevalent (but variably so across sites). Moreover, *P. leucopus* is the most common small mammal at all eight locations (Fig. 1A). Samples sizes for this study are: 1) **UNDE** - University of Notre Dame Environmental Research Center, Michigan (*N* = 22, 9♂ 13♀). 2) **STEI** - Steigerwalt-Chequamegon, Wisconsin (*N* = 27, 15♂ 12♀). 3) **HARV** - Harvard Experimental Forest & Quabbin Watershed, Massachusetts (*N* = 36, 23♂ 13♀). 4) **MLBS** - Mountain Lake Biological Station, Virginia (*N* = 21, 14♂ 7♀). 5) **SCBI** - Smithsonian Conservation Biology Institute, Virginia (*N* = 17, 9♂ 8♀). 6) **BLAN** - Blandy Experimental Farm, Virginia (*N* = 12, 9♂ 3♀). 7) **SERC** - Smithsonian Environmental Research Center, Maryland (*N* = 8, 6♂ 2♀). 8) **ORNL** - Oak Ridge National Laboratory, Tennessee (*N* = 5, 2♂ 3♀). Animal collections were performed under authorization from IACUC (# IS00009477).

### Climate data

We downloaded 2022 climate data from the NEON data portal (https://www.neonscience.org/data) and then used these data to describe climatic variation at each site. We first calculated total annual precipitation (mm), mean annual relative humidity (%), and mean annual temperature (°C) measured at thirty-minute intervals over the year (data products DP1.00006.001 and DP1.00098.001) for each of the eight sites. Then, we calculated the monthly total precipitation values, mean temperature, and mean relative humidity to calculate coefficients of variation (CV) for each of the three climate variables and used all six variables (3 annual means and 3 monthly CVs) in a principal components analysis (unpublished data). We used PC1 from this PCA as the ‘climate’ variable in our gene expression analyses.

### Molecular data

#### Target gene sequences and primer and probe design

Candidate gene sequences for ddPCR (droplet digital PCR) assays were first identified within the *P. leucopus* or *P. maniculatus* genomes using GenBank. Then, primers and probes (Supplementary material, Tables S2 and S3) for all six genes were designed using the Primer Quest online tool from Integrated DNA Technologies (IDT), using qPCR parameters (2 primers + probe). ZEN double-quenched probes were then chosen, with either a FAM or HEX fluorescent dye used for different genes. Compared to TaqMan probes, ZEN probes are preferred for ddPCR due to higher discrimination between positive and negative droplets (see below). For assays, all primers and probes were diluted to 10 µM concentration. To ensure the primers and probes functioned as intended, a synthetic DNA (sDNA) strand of each target sequence was created using IDT’s g-Block gene fragment product (Supplementary material, Table S4). Then, various primer/probe, sDNA, and complementary DNA (cDNA) from *P. leucopus* concentrations were investigated to optimize ddPCR assays, both as a single-plex for each gene and for gene triplexes A and B (Supplementary material, Tables S5-S7 and Fig. S1-S3).

#### RNA extraction

Ear biopsies (< 1mg) were transferred to 2mL reinforced screwcap tubes (Fisherbrand™ Bulk tubes, 15-340-162) with 2 ceramic beads (2.8mm; Fisherbrand™ Bulk beads, 15-340-160) and 500 µl cold (4°C) Trizol reagent (Invitrogen, 15596018). Samples were then homogenized (Fisherbrand™ Bead Mill 24 Homogenizer, 15-340-163) at 4.5m/s for 5 cycles of 30 seconds with 30-second intervals, and homogenates were transferred to sterile 1.5 mL microtubes and stored for 5 min at room temperature (23°C) to allow the dissociation of nucleoprotein complexes. Cold chloroform (100 µl) was then added to tubes (4°C), vortexing vigorously for 15 seconds. This mixture was stored for 5 min at room temperature and then centrifuged (12,000 x g, 15 min, 4°C). The clear layer containing RNA was transferred to a new 1.5 mL microtube. In these new microtubes, 250 µl cold (4°C) isopropyl alcohol was added to precipitate RNA with 1.25 µl of 20mg/ml of glycogen (RNA grade, Thermo Scientific, R0551) to improve pellet visualization. Samples were then agitated (10 sec) and incubated overnight in a −20°C freezer. The following day, samples were centrifuged (12,000 x g, 10 min, 4°C), supernatants were disposed, and 500 µl cold ethanol (75%, 4°C) was added to each microtube to wash samples. The samples were then centrifuged again (12,000 x g, 5 min, 4°C), the supernatants disposed, the samples centrifuged once more (12,000 x g, 3 min, 4°C), and the remaining ethanol evaporated. The dried pellets were then resuspended with 30 µl RNase-free water, and due to the small size of the ear samples and the low RNA yields, RNA concentration of all samples was measured via Qubit (Invitrogen, Qubit 4 Fluorometer, Thermo Scientific, USA). Quantification of genomic DNA (gDNA) was also performed for a subset of samples, and a general linear model (GLM) analysis revealed that gDNA concentration effects on gene expression were non-significant (F_(5, 54)_ ≥ 0.361, *p* ≥ 0.104; Supplementary material, Table S8).

### Conversion of RNA into complementary DNA (cDNA)

Reverse transcription was performed for each extracted RNA sample using 500 ng RNA, 4 µl 5X reaction mix, and 1 µl reverse transcriptase (iScript cDNA Synthesis kit, 1708891, Bio-Rad) in RNase-free water to obtain a total volume of 20 µl, following the manufacturer’s instructions. Samples were placed in a thermocycler, following the program: 25°C for 5 min, 46°C for 20 min, 96°C for 1 min, and holding at 4°C.

### Droplet Digital Polymerase Chain Reaction (ddPCR)

Droplet digital PCR uses a water-in-oil micro-droplet approach to quantify absolute copy numbers of a PCR target (Hindson et al. 2011; Whale et al. 2016). Given our obligate reliance on ear tissue, which is modestly bioactive, ddPCR was the ideal technology Hindson *et al*. 2011; Whale *et al*. 2016). Droplet digital PCR reactions contained 5µl ddPCR Multiplex Supermix (ddPCR Multiplex Supermix, 12005909, Bio-Rad); 4.5 µl target primer (10 µM; forward + reverse mix), 0.63 µl probe FAM, 0.63 µl probe HEX, and 0.63 µl probe FAM + HEX (e.g., when 50% FAM + HEX, add 0.31 µl of each; when 60% FAM + 40% HEX, add 0.38 FAM + 0.25 HEX), and 6 µl sample (cDNA 25ng/µl). Next, 22 µl of each sample was gently transferred to the middle row of the cartridge (DG8™ Cartridges and Gaskets, 1864007, Bio-Rad), and 71 µl of droplet generation oil (Droplet Generation Oil for Probes, 863005, Bio-Rad) was added into the bottom row of the cartridge. Cartridges were covered with gaskets and placed on a droplet generator (QX200™ Droplet Generator, 1864002, Bio-Rad). Each sample was apportioned into 15,000-20,000 nanoliter-sized oil droplets, which then were transferred to a 96-well plate (ddPCR™ 96-Well Plates, 12001925, Bio-Rad).

Plates were covered with foil (PCR Plate Heat Seal, foil, pierceable 1814040, Bio-Rad), sealed (PX1 PCR Plate Sealer, 1814000, Bio-Rad), and transferred to a thermal cycler (C1000 Touch™ Thermal Cycler with 96–Deep Well Reaction Module, 1851197, Bio-Rad). Reactions conditions were: 95°C for 10 min, followed by 48 cycles of 94°C for 50 sec and 58°C (Triplex A) or 62°C (Triplex B) for 2 min. These cycles were followed by a melting curve of 98°C for 10 min and holding at 4°C. After amplification, the droplets were separated and counted as either positive (i.e., having the target sequence of interest) or negative (i.e., not having the target sequence of interest) (Taylor et al. 2017) using the droplet reader (QXDx Droplet Reader, 12008020, Bio-Rad). At the end of all runs, expression data were obtained using QuantaSoft^TM^ Analysis Pro software (version 1.05).

### Statistical Analyses

Our primary interest was to reveal whether and how climate conditions at NEON sites related to the competence of individual mice for *B. burgdorferi*, not how climate affected single genes. Subsequently, we conducted a Varimax-normalized principal component analysis (PCA) on log_10_ transformed gene expression data to discern whether and how genes were expressed as a collective within individual mice. Following the Kaiser criterion, we considered only principal components with eigenvalues > 1 as appropriate for further investigation. This PCA revealed two PCs with eigenvalues > 1, which we termed ‘resistance’ and ‘tolerance’ based on their genetic composition (Table S1).

To discern how climate and other forces affected variation in gene expression (PC1 and PC2), we followed a model-fitting procedure. We compared AICc scores of a series of progressively simpler models, starting from an omnibus model, including mouse sex (male or female), mouse body mass, and climate at a site (climatic PC score), and all possible interactions as fixed effects. Tick presence (presence or absence on an individual mouse) was also included as a simple fixed effect, but no interactions for this term were included as we lacked tick data for many mice (*N* = 54 of 148 total mice lacking tick data). We also included NEON sites as a random effect in all models to account for climate being a site-level variable.

## Results

Our first goal was to discern whether and how immune gene expression clustered among individual mice (*N* = 148) across all eight NEON sites. For this, we used principal component analysis (PCA), which recovered two dimensions of variation in gene expression, generally representing *B. burgdorferi* resistance and tolerance. Specifically, PCA revealed two components with eigenvalues greater than 1, accounting for 87.21% of the total variation in gene expression among individuals (Table S1A). The first component (PC1, 55.01% variation) was positively associated with *TLR2*, *IFN-γ*, *IL-6*, and *IL-10* expression (Table S1B), PC scores for which we termed resistance. The second component (PC2, 32.20%) was positively related to TGF-β and GATA3 expression (Table S1B), PC scores for which we termed tolerance.

We used linear mixed models to analyze resistance and tolerance PC scores separately, starting with the same omnibus model structure. Each omnibus model included the main effects of individual mouse sex, individual body mass, climate conditions at the NEON site (unpublished data), and all possible two and three-way interactions among these variables. NEON site was also always included as a random term to accommodate the non-independence of individuals and climatic conditions at the same site. We also included a simple binary indicator of tick presence, but we did not include interaction terms with tick presence in our models due to insufficient coverage for this variable. NEON also did not have access to a higher-resolution estimate of tick burden (i.e., count or score) for these mice, so this binary variable was the only option we had to discern vector effects on host immune gene expression. Starting with omnibus models, we identified best-fit models for resistance and tolerance separately by systematically removing terms from more complex models until we minimized Akaike information criteria scores for small sample sizes (AICc) (Table 1). For resistance, the best-fit model included only climate. This model explained a large amount of the variation either without (conditional R^2^ = 31%) or with (marginal R^2^ = 28%) the random effect of NEON sites. Mice from warmer, wetter sites exhibited greater expression of resistance to *B. burgdorferi* (Fig. 1B). By contrast, climate was not a strong predictor of tolerance. The best-fit model for tolerance included only individual sex (Table 1), but the disparity in the variance explained by the marginal R^2^ and conditional R^2^ values suggest that the effect of sex on tolerance might vary among sites but be difficult to detect here because of the over-representation of some sexes in some NEON sites. Moreover, model weights (*w*) for the top 4-5 models for tolerance overlap a bit more than for resistance; indeed, the best-fit model for resistance is >3x more explanatory than the next best model. Altogether, these results suggest that whereas other factors we considered for tolerance do have minor predictive power, none of these relationships is nearly as strong and obvious as climate effects on resistance.

**Table 1.**
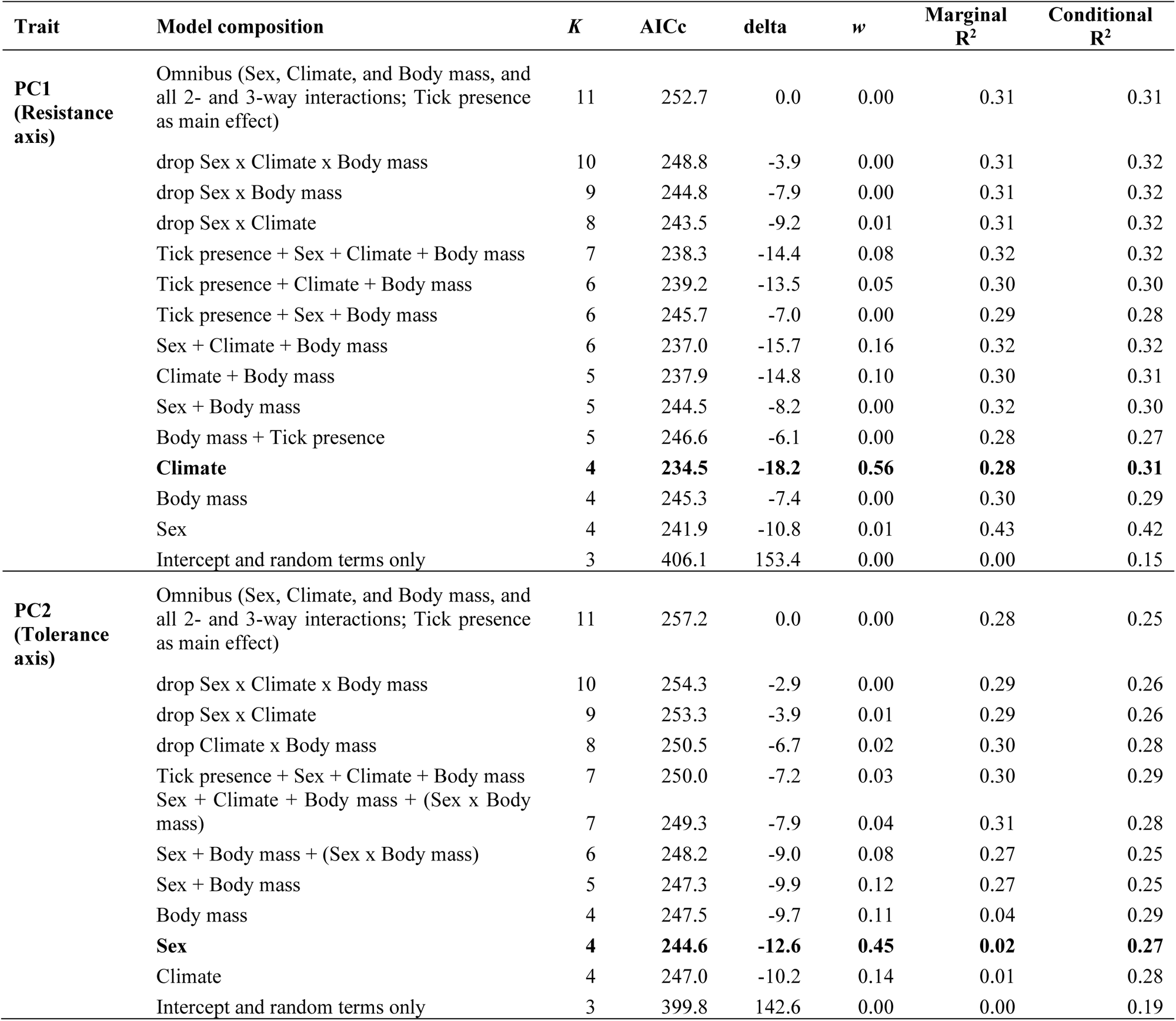
Comparisons of models to predict immune gene expression in *Peromyscus leucopus* ear tissue biopsies.

## Discussion

*P. leucopus* from eight locations, spread across much of the northeastern U.S., differed in expression of immune genes integral to protection against *B. burgdorferi* infection. The dominant aspect of immune gene variation among individuals comprised a putative resistance axis, a continuum of expression of TLR2, IFN-γ, IL-6, and IL-10. A second orthogonal dimension of variation involved the expression of TGF-β and GATA3, a putative tolerance axis. Most intriguingly, *P. leucopus* from warmer and wetter NEON sites had, on average, more resistant gene expression profiles than mice from colder and drier areas (Fig. 1B). Tolerance was comparatively unexplained by the variables we considered. Below, we first discuss whether and how our simple immune gene measures might relate to the complex processes by which this host interacts with *B. burgdorferi* and the black-legged ticks that transmit it. We then discuss how host populations could come to vary in their immunological disposition, and we summarize possible ecological consequences for this variation in *B. burgdorferi* risk across the landscape.

### Immunity to B. burgdorferi in rodents

Initial vertebrate responses to *B. burgdorferi* exposure involve a variety of rapid, innate immune activities essential to the establishment of an active infection. The first lines of defense include phagocytosis by macrophages and neutrophils, and oxidative burst and antimicrobial peptide activity by various leukocytes (Montgomery and Malawista 1996). *Borrelia burgdorferi* molecular motifs are first recognized by TLR2, initiating a local inflammatory response including cytokines such as IFN-γ, which activates some local cytotoxic T cells and macrophages, and IL-6, which drives the differentiation of B cells and antibody responses, all leading to pathogen clearance (Akdis et al. 2016; Kopitar-Jerala 2017; Bockenstedt et al. 2022). Conversely to these factors, IL-10 plays an important role in limiting immunopathology (Iyer and Cheng 2012), which is partly accomplished by inhibiting activity of natural killer (NK) cells and deactivating macrophages. What is particularly relevant for understanding *P. leucopus* as a *B. burgdorferi* reservoir, though, is that *P. leucopus* (relative to lab model organisms and other putative hosts of *B. burgdorferi*) endure high *B. burgdorferi* burdens for sometimes long periods without overt signs of disease (Barbour 2017). Ecologically, this trait is important, as nymphal and larval ticks in the Northeastern U.S. are strongly, seasonally asynchronous (Ogden et al. 2021). *B. burgdorferi* must, therefore, spend a considerable amount of time in a vertebrate host (sometimes several months) before transmission to the next generation of larval ticks is possible (Voordouw et al. 2015). Immunologically, such a type of tolerance of *B. burgdorferi* infection by *P. leucopus* is probably due to sustained but mild inflammation (Bourgeois et al. 2024). Once activated, macrophages in *P. leucopus* tend to adopt less inflammatory phenotypes than those of *Mus musculus* (Barbour 2017; Balderrama-Gutierrez et al. 2021; Milovic et al. 2024). By contrast, *P. leucopus* neutrophils become more active and prone to degranulation upon *B. burgdorferi* exposure, but collateral damage from these processes is partly ameliorated in *P. leucopus* by secretory leukocyte peptidase inhibitor (*Slpi*) and superoxide dismutase 2 (*Sod2*; (Milovic et al. 2024)). Perhaps the most novel means by which *P. leucopus* (and other species in this genus) avoid disease when infected with *B. burgdorferi* is the lack of a high-affinity Fc immunoglobulin gamma receptor I (FcγRI) or CD64 (Barbour et al. 2023). These factors also make sustained inflammatory responses comparatively mild (Ioan-Facsinay et al. 2002). Presently, it is obscure how the six specific genes we measured relate to these various mechanisms, but we chose these six genes because they are implicated in the control of *B. burgdorferi* in many vertebrates, including *P. leucopus* (Long et al. 2019). Additional higher-resolution data on the protective roles of the factors we measured in pinnae skin would be valuable here.

### Causes of immunological variation among Peromyscus leucopus

We recognize the importance of knowing if the resistance immune profile we found directly translates into infection intensity – unfortunately, we can’t do that with our study – this is not a lab study, and it was done over an area spanning hundreds of miles, we have no way to know when/if these mice were exposed to *B. burgdorferi.* Moreover, recent exposure would have profoundly different effects on immune variation than old exposures. Currently, there is no way we could ascertain *Borrelia*’s infection status, so the best we could do was to determine whether a tick was on the animal. Even in controlled lab experiments, measuring *Borrelia* in skin biopsies has been challenging. According to Bourgeois et al. (2024), it is possible to measure *Borrelia* burden using ddPCR in the ears of *Mus musculus* mice (C57BL/6J and C3H/HeN) after two-weeks post-infection; however, the same infection done in *Peromyscus leucopus* mice was only detected after one month.

Additionally, considering the scale of our study, the immunological variation we described among sites probably arises through a mix of genetic (G), environmental (E), and GxE interactive factors (Martin et al. 2021). Since we measured six genes with unique evolutionary history, it is reasonable to expect some genetic differences among our eight focal host populations. However, we expect that much, if not most, variation in immune gene expression to be derived from phenotypic plasticity. Rodent immune systems are highly sensitive to environmental conditions (Martin et al. 2021), particularly food (Thomason et al. 2013; Blubaugh et al. 2023). For example, in *P. maniculatus*, mild food restriction compromised the ability of individuals to mount antibody responses to a non-pathogenic antigen (Derting and Compton 2003; Martin et al. 2007).

We expect the immune variation we observed to be at least partly attributable to resource availability among sites. Climate conditions across sites, and especially over the years, affect oak mast and other food resources (Jones et al. 1998; Ostfeld et al. 2018), which should alter both the ability of mice to build and mobilize defenses and the need to forage. Of course, we also cannot discount more complex effects of environmental conditions on *B. burgdorferi* ecology, including mesopredator release (Ostfeld et al. 2018), which could affect *P. leucopus* behavior, immunity, stress physiology, or survival (Levi et al. 2012). We also cannot rule out allelic differences among populations that could underpin some variation in gene expression. Indeed, a recent study in coral found that white band disease prevalence and susceptibility increased with temperature stress, but resistance to the disease was related to genetic variation in four immune genes among populations (Vollmer et al. 2023). Regardless of the specific combination of factors, some interplay of environments and genes underlie an appreciable level of variation in *B. burgdorferi* resistance in *P. leucopus* (Plowright et al. 2024).

### Conclusions, limitations, and future directions

Historically, epidemiological models have taken simplifying approaches to understand species’ contributions to multi-host infectious diseases (Anderson and May 1979). Now though, we know that some individual hosts within species are disproportionately more influential than others in many disease systems (Martin et al. 2019). We worry that these understandable simplifications might have unintentionally led to idealistic assumptions in empirical realms of infectious disease biology, such as the search for zoonotic hosts, including SARS-COV2 (Griffin et al. 2021). For instance, *Peromyscus leucopus* and ∼30 other mammalian species (Jemeršić et al. 2021) have been implicated as points of vulnerability in the management of COVID risk (Fischhoff et al. 2021), but our data suggest that for the most common zoonotic pathogen in North America, the same host species can serve as a risk source or sink, depending on context. We expect the same holds for many, if not most, zoonoses. One of the most surprising results of our study was how little a role mouse sex seemed to have for gene expression differences. In nature, male *P. leucopus* often have higher tick burdens than females (Brunner and Ostfeld 2008), and *B. burgdorferi* infection can even alter the rates at which males are recaptured (Voordouw et al. 2015), a particularly surprising effect (given the larger home range size of males than females).

We still think that the large-scale pattern is very interesting and worthwhile, and it’s difficult to contest that there is a cool/significant pattern. Our study, like all studies, ultimately ends with many questions and future directions – we don’t view it as a critical flaw that we don’t know everything about resistance at geographical scales – what we find most intriguing is that there’s a pattern we’ve uncovered (even if it could be driven, in whole or part, by many things we don’t study). Thus, our study opens new possibilities to consider when studying disease dynamics. Given the complexity of the *B. burgdorferi* system, the amenability of NSF’s NEON to large scale studies like ours, and the importance of understanding Lyme disease ecology for human health, we encourage additional integrative approaches to future zoonosis surveillance and mitigation, efforts informed by appropriate degrees of ecological, evolutionary, and public health perspectives.

## Acknowledgments

We thank the NEON technicians, particularly Ashley Spink, Josh Fischer, Lindsey Hayter, and Will Ponder, for collecting the ear samples for our research. Sara Paull, Samantha Kremidas, and Laura Steger from NEON/Battelle for their flexibility in adapting protocols to meet our project’s needs. We are also grateful to Dr. Maria Diuk-Wasser of Columbia University for generously providing lab-raised *Peromyscus leucopus* ear samples, which were instrumental in developing our methods. Additionally, we wish to acknowledge the significant contributions of our undergraduate assistants, Gabby Mansilla and Kevin Galassini, as well as Ph.D. student D. Tucker Simonton, for their help collecting the molecular data.

## Funding

National Science Foundation grant 2110070 (LBM, JLO).

## Author Contributions

Conceptualization: VRA, JLO, LBM. Methodology: VRA, KM, RAM. Data collection: VRA, AMB. Data analyses: VRA, AMB, LBM. Funding acquisition: LBM, JLO. Writing – original draft: VRA, LBM. Writing – review & editing: VRA, KM, RAM, JLO, LBM.

## Competing Interest Statement

Authors declare that they have no competing interests.

## Notes

### Competing Interest Statement

The authors have declared no competing interest.

