## Supplementary material for "Climate drives geographic variation in individual *Peromyscus leucopus* immunity against zoonotic disease"

**This file includes:**

3 Figures: S1-S3

8 Tables: S1-S8

Two droplet digital PCR (ddPCR) multiplexes were developed to measure the expression of six distinct immune genes in *P. leucopus* pinnae biopsies: **Multiplex A:** IFN-γ, IL-6, and IL-10. **Multiplex B:** TLR2, GATA3, and TGF-β.

**Gene selection**

We selected genes that play strong and distinct roles in the host defense against *Borrelia burgdorferi* infection. These genes include pattern recognition receptor for the bacteria (TLR2) and cytokines that foster resistance (IFN-γ and IL-6) and tolerance (IL-10, TGF-β, and GATA3) (Table S2).

**Primers and probes design**

Primers and probes (Table S3) were designed using the Primer Quest online tool from Integrated DNA Technologies (IDT), using qPCR parameters (2 primers + probe). ZEN double-quenched probes were then used with a FAM or HEX fluorescent dye. When deciding which genes should receive a FAM or HEX probe, Bio-Rad recommends that your main target of interest should be measured using the FAM fluorescence because of the better discrimination on the ddPCR interface.

**Synthetic DNA**

To discern how little gene expression we could measure in the absence of artifacts arising in the extraction or cDNA synthesis steps, we obtained synthetic DNA (sDNA) of each target sequence (i.e., IDT's g-Block). Each g-Block was between 125-250bp (Table S4). All sDNA arrived in lyophilized form but were reconstituted using 25µl Tris-EDTA buffer before assays to 10ng/µl. These sDNA samples were further diluted to 0.0001 ng/µl for ddPCR validation.

**Development of the droplet digital PCR (ddPCR) assays with synthetic DNA (sDNA)**

A droplet digital PCR (ddPCR) single-plex was first performed for each gene using the appropriate synthetic DNA (sDNA) to ensure the primers and probes amplified the target of interest. The first step was to create a reaction containing the following materials: ddPCR Supermix for probes (Cat. No. 186-3010, Bio-Rad), primers, probes, nuclease-free water, and 5ul of sDNA (Table S5). The reaction mix (25ul) was loaded on an eight-channel droplet generator cartridge (Cat. No. 186-3008, Bio-Rad). The droplets were then generated with 70ul of droplet generation oil (Cat. No. 186-3005, Bio-Rad) in the droplet generator of the QX100 system (Bio-Rad). The generated droplets were carefully transferred to a 96-well PCR plate. The PCR plate was then thermal sealed with pierceable foils in a PCR plate sealer PX1 (Cat. No. 181-4040) and placed into the PCR machine. The standard PCR settings recommended by Bio-Rad were used for the initial ddPCR runs. After the PCR cycle, the sealed plates were placed in the droplet reader, and the plate was analyzed according to the manufacturer's recommendations. Next, both triplexes were run using sDNA with the same PCR parameters as above. The only difference was that the amount of sDNA added to the reaction included all three targets. Each single-plex revealed amplification of only our target sequences (Fig. S1). We also found that adding non-target sDNA to each single-plex did not yield any amplification; therefore, the primers and probes only amplified the target of interest. Each multiplex also revealed amplification of only target sDNA; no non-target amplification was detected for any gene in either triplex (Fig

S1). We also found that when we changed the amount of sDNA input into reactions (increments of 0.5ul, ranging from 1 – 4ul), gene expression increased linearly as expected for both triplexes (Fig. S2).

**Optimizing ddPCR assays with *Peromyscus leucopus* ear biopsies**

*Peromyscus leucopus* ear tissues (immediately flash-frozen upon collection) were obtained from Maria Diuk-Wasser for further assay optimization. RNA was extracted, and cDNA was synthesized from these tissues, as described above, for samples in the main study. To ensure assays involving cDNA from *P. leucopus* performed as well as in ddPCR as sDNA samples, both triplexes were run on the same samples at varying levels of cDNA input (i.e., 1ul, 2ul, 3ul, and 4ul). Then, to determine the optimal concentration of cDNA that would yield high repeatability and also assess whether assay results would vary among runs or researchers performing assays, samples were transcribed to 50ng, 100ng, and 150ng and run by three different people in triplicate. For both triplexes, ddPCR methods for sDNA were effective for *P. leucopus* cDNA. No further optimization of temperatures, cycles, or primer/probe concentrations was necessary (Fig. S3). Interpersonal variation was also low, and repeatability across runs was high. For replicate assay runs using 150ng cDNA, CVs for each gene in triplex A was < 10%; for triplex B, all CVs < 20% (Table S6). Across researchers (same researcher, 4 assays), expression measured in both triplexes had CVs < 12% (Table S7).

As ear biopsies available from NEON were so small, we next sought to rule out the possibility that gDNA might confound our expression measurements. Indeed, total RNA yields of pinnae samples tended to be low, and DNase treatment was inadvisable.

Subsequently, to discern whether gDNA contamination might affect expression outcomes, we quantified both RNA and gDNA using Qubit in a subset of our samples (60 out of 148 total samples = 40%), then used a General Linear Model (GLM) with gene expression as the dependent variable and gDNA contamination, which was calculated as (([gDNA] / [RNA]) * 100) as a predictor. As shown in Table S8, gDNA contamination did not significantly affect gene expression.

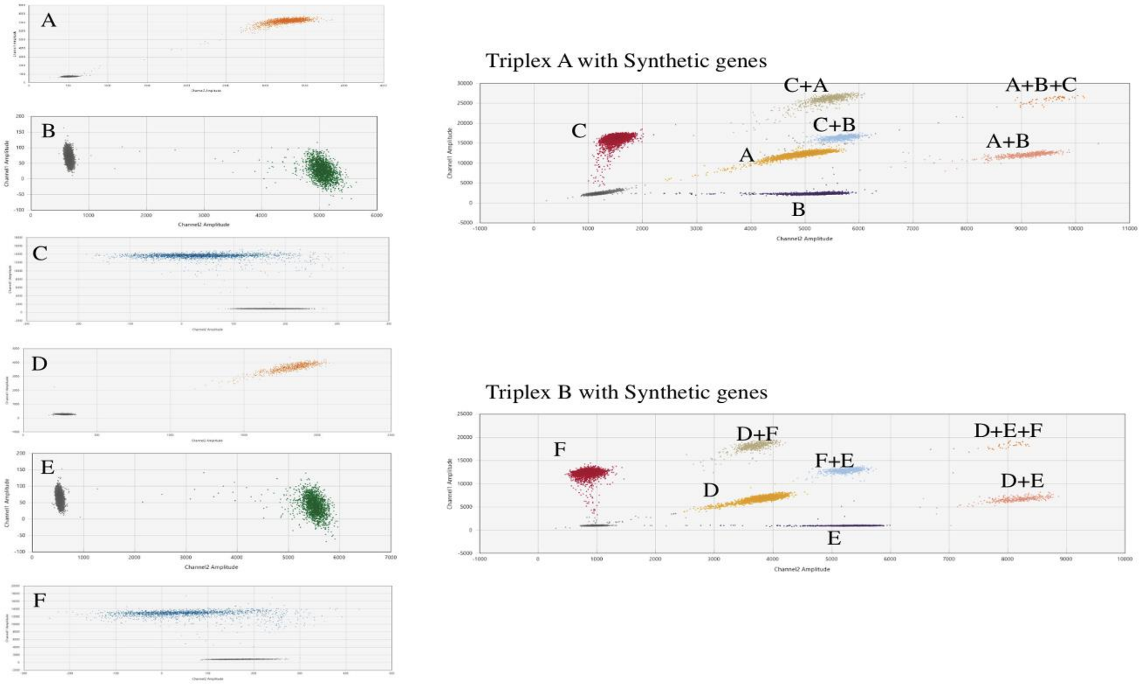


**Fig. S1.** Synthetic DNA (sDNA) single-plexes and multiplexes 2D results. **A**: IL-6, **B**: IFN-γ, **C**: IL-10, **D**: GATA3, **E**: TLR2, and **F**: TGF-β.

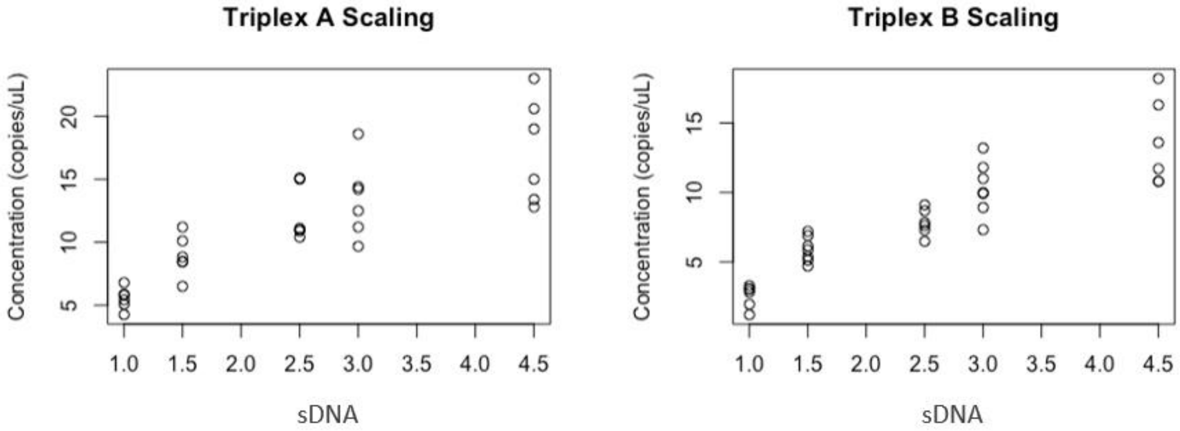


**Fig. S2.** Multiplex performance is consistent across sDNA concentrations. With the increments of 0.5ul (1 - 4ul) in the sDNA volume, we see an increase in the gene expression.

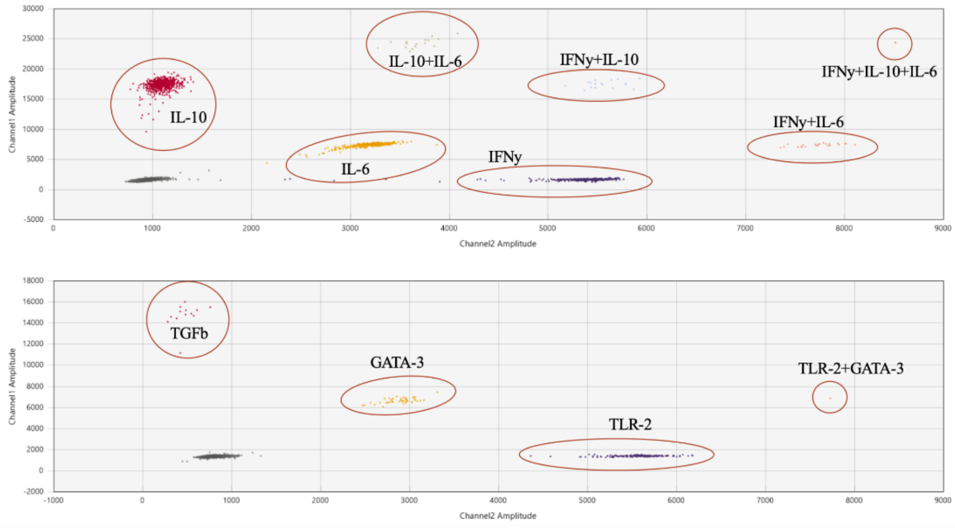


**Fig. S3.** Example or one run of optimized ddPCR results using cDNA synthesized from RNA extracted from pinnae of *Peromyscus leucopus*. The upper plot shows triplex A, and the lower plot shows triplex B.

**Table S1A.** Components retained by principal component analysis (PCA) of six immune gene targets among all *Peromyscus leucopus*.

| Total Variance Explained | | | |
|---|---|---|---|
| | Initial Eigenvalues | | |
| **Component** | **Total** | **% of Variance** | **Cumulative %** |
| **1** | 3.301 | 55.01 | 55.01 |
| **2** | 1.932 | 32.20 | 87.21 |
| **3** | 0.384 | 6.40 | 93.61 |
| **4** | 0.296 | 4.93 | 98.53 |
| **5** | 0.053 | 0.88 | 99.41 |
| **6** | 0.035 | 0.59 | 100.00 |

Extraction Method: Principal Component Analysis.

**Table S1B.** Individual immune gene expression loadings on each of the two PCA axis (after varimax rotation with Kaiser normalization).

| | Component | |
|---|---|---|
| | **1** | **2** |
| **TLR2** | **0.850** | -0.025 |
| **IFN-γ** | **0.857** | 0.016 |
| **IL-6** | **0.965** | -0.065 |
| **IL-10** | **0.944** | -0.037 |
| **TGF-β** | -0.050 | **0.985** |
| **GATA3** | -0.007 | **0.987** |

**Table S2.** The defensive roles of each of the six immune genes studied here.

| Targets | Function |
|---|---|
| Toll-like receptor 2 **(TLR2)** | Recognizing the proteins of the spirochete bacteria of the *Borrelia burgdorferi sensu lato* |
| Interferon-gamma **(IFN-y)** | Main mediator of the effector mechanisms of a Th1 response |
| Interleukin 6 **(IL-6)** | Responds to tissue damage and has both pro and anti-inflammatory properties |
| Interleukin 10 **(IL-10)** | Inhibits inflammatory cytokines produced by macrophages in response to *B. burgdorferi* |
| Transforming growth factor β **(TGF-β)** | Plays an important role in wound healing and has anti-inflammatory functions |
| GATA binding protein 3 **(GATA3)** | A mediator of Th2 response and determined to be an immunological maker of tolerance to macroparasites in voles |

Th1 response: i.e., activation of cytotoxic T cells and macrophages
Th2 response: i.e., differentiation of B cells and antibody production

**Table S3.** Primer and probe sequences for each target gene

| Multiplex | Gene | Sequence (5' → 3') |
|---|---|---|
| A | IFN-y | **F:** CTG TGG GTG GTA ACC GAT TT |
| | | **R:** CGT TGG CTC AGG TTG TAG AT |
| | | **Probe:** /5HEX/ACT ACA GTG /ZEN/ATG CCT TGA GCT GCT /3IABkFQ/ |
| | IL-6 | **F:** CTT CCA TCC ACT TGC CTT CT |
| | | **R:** TCC TCT GTG AAG TCT CCT CTC |
| | | **Probe:** /56-FAM/CCA CTG CCC /ZEN/TTC CTA CCT CAC AAG /3IABkFQ/ |
| | | **Probe-** /5HEX/CCA CTG CCC /ZEN/TTC CTA CCT CAC AAG /3IABkFQ/ |
| | IL-10 | **F:** TGG GAA GCC AAC TGA AGC |
| | | **R:** CTC TGA ACC CAG GAA GGA AAG |
| | | **Probe:** /56-FAM/ACA CCA CAG /ZEN/TAA ACA CGT CGG TAG C/3IABkFQ/ |
| B | TLR2 | **F:** CCT GTT GAT CCT GCT CAT AGT C |
| | | **R:** TTT CTT GGG CTT CCT CTT GG |
| | | **Probe:** /5HEX/TAC CTG AGA /ZEN/ATG ATG TGG GCG TGG /3IABkFQ/ |
| | TGF-β | **F:** CTG AAC CAA GGA GAC GGA ATA C |
| | | **R:** GGG ACT GAT CCC GTT GAT TT |
| | | **Probe:** /56-FAM/TTC AGC GCT /ZEN/CAC TGC TCT TGT GA/3IABkFQ/ |
| | GATA-3 | **F:** CTA CTG GGT TCG GGT GTA AG |
| | | **R:** GTA GTG CCC AGT ACC ATC TC |
| | | **Probe:** /56-FAM/AGG CAG GGA /ZEN/GTG TGT GAA CTG TG/3IABkFQ/ |
| | | **Probe:** /5HEX/AGG CAG GGA /ZEN/GTG TGT GAA CTG TG/3IABkFQ/ |

Abbreviations are as follows: **F:** forward primer; **R:** reverse primer; **HEX:** hexachlorofluorescein; **FAM:** fluorescein amidite.

**Table S4.** G-blocks, single-stranded synthetic DNA (sDNA) sequences for each target gene

| Gene | Sequence (5’ → 3’) |
|---|---|
| **TLR2** | CCCTCAGTCTTGGAGTGCCACCAGGCTCTACTAGTGTCTGGCGTCTGCT GTGCCCTTCTCCTGTTGATCCTGCTCATAGTCGGCCTGTGCCACCATTTC CACGGGCTGTGGTACCTGAGAATGATGTGGGCGTGGCTCCAGGCCAAG AGGAAGCCCAAGAAAGCTCCATGCAGGGACATTTGCTATGATGCCT |
| **IFN-y** | AGGGGCCAGTCAACCAGTTGACTGAAGTCAGACTGTGGGTGGTAACCG ATTTTTACTTGACAATGAGGAACACTCACTACAGTGATGCCTTGAGCTGC TGCTGGCCGGAGAGCGCATCTACAACCTGAGCCAACGCCTCAATAGCCG GTCAGCACTGTGACGATGCATC |
| **IL-6** | CTATGAAGTTCCTCTCCGCAAGAGACTTCCATCCACTTGCCTTCTTGGGC CTGTTGCTGGCGATGGCCACTGCCCTTCCTACCTCACAAGTCCGGAGAG GAGACTTCACAGAGGACACCACTCCCAACAGACC AGTGTAT |
| **IL-10** | ATATTCTATAATGGGGTGGGGGAGGGGGTCTTCTTTGGGAAGCCAACTG AAGCTTCCGTTCTAAGGCTGGCCACACTTTATAGCTACCGACGTGTTTAC TGTGGTGTTCTCTAATTTCTTTCCTTCCTGGGTTCAGAGCTCCTGACGTA GTTGTGAAGACTCTTACAGGA |
| **TGF-β** | CGTCACCGCAGTCGTACGGCAGTGGCTGAACCAAGGAGACGGAATACA GGGCTTTCGCTTCAGCGCTCACTGCTCTTGTGACAGCAAAGATAACGTA CTCCACGTGGAAATCAACGGGATCAGTCCCAAACGTC GAGGTGACCTGGGCACCATTCATGACAT |
| **GATA3** | ACTCTTCCCACCCAGCAGCCTGCTGGGAGGATCCCCTACTGGGTTCGGG TGTAAGTCGAGGCCCAAGGCGCGGTCCAGCACAGGCAGGGAGTGTGTG AACTGTGGGGCCACCTCCACCCCACTGTGGCGGCGAGATGGTACTGGG CACTACCTCTGCAACGCCTGCGGACTCTACCATAAGAT |

**Table S5:** ddPCR reaction setup per well

| **Component** | **Volume per 25ul Reaction** |
| --- | --- |
| Supermix | 5.00 |
| Forward Primer (10uM) | 2.25 |
| Reverse Primer (10uM) | 2.25 |
| Probe (10uM) | 0.63 |
| RNase free water | 9.87 |
| sDNA | 5.00 |
| Total | 25.00 |

**Table S6.** Interpersonal variation experiment, including the standard deviation for all genes, mean, and corresponding coefficients of variation.

| cDNA concentration | Target | Standard deviation | Mean | Coefficient of variation |
|---|---|---|---|---|
| **50ng** | IL-10 | 1.088 | 9.16 | 12% |
| | IFN-γ | 1.389 | 7.575 | 18% |
| | IL-6 | 1.177 | 7.721 | 15% |
| | TGF-β | 0.428 | 0.62 | 69% |
| | GATA3 | 0.028 | 0.442 | 6% |
| | TLR2 | 1.711 | 6.18 | 28% |
| **100ng** | IL-10 | 4.15 | 20.075 | 21% |
| | IFN-γ | 2.638 | 15.525 | 17% |
| | IL-6 | 3.761 | 15.249 | 25% |
| | TGF-β | 0.214 | 4.415 | 5% |
| | GATA3 | 0.615 | 4.415 | 14% |
| | TLR2 | 1.697 | 14.1 | 12% |
| **150ng** | IL-10 | 2.95 | 30.763 | 10% |
| | IFN-γ | 2.136 | 24.45 | 9% |
| | IL-6 | 2.354 | 24.8 | 9% |
| | TGF-β | 0.021 | 0.746 | 3% |
| | GATA3 | 0.53 | 6.145 | 9% |
| | TLR2 | 0.707 | 21.9 | 3% |

**Table S7.** Intrapersonal variation experiment, including the standard deviation for all genes, mean, and corresponding coefficient of variation.

| cDNA concentration | Target | Standard Deviation | Mean | Coefficient of Variation |
|---|---|---|---|---|
| **50 ng** | IL-10 | 1.09 | 9.16 | 12% |
| | IFN-γ | 1.39 | 7.58 | 18% |
| | IL-6 | 1.18 | 7.72 | 15% |
| | TGF-β | 0.227 | 0.519 | 44% |
| | GATA3 | 0.975 | 1.796 | 54% |
| | TLR2 | 4.543 | 8.163 | 56% |
| **100 ng** | IL-10 | 4.15 | 20.08 | 21% |
| | IFN-γ | 2.64 | 15.53 | 17% |
| | IL-6 | 3.76 | 15.25 | 25% |
| | TGF-β | 0.399 | 0.907 | 44% |
| | GATA3 | 1.619 | 4.583 | 35% |
| | TLR2 | 1.817 | 14.333 | 13% |
| **150 ng** | IL-10 | 2.95 | 30.76 | 10% |
| | IFN-γ | 2.14 | 24.45 | 9% |
| | IL-6 | 2.35 | 24.8 | 9% |
| | TGF-β | 0.161 | 0.944 | 17% |
| | GATA3 | 1.317 | 7.02 | 19% |
| | TLR2 | 3.966 | 22.243 | 18% |

**Table S8.** Residual gDNA in pinnae extracts was not related to gene expression.

| Tests of Between-Subjects Effects | | | | | | | |
|---|---|---|---|---|---|---|---|
| **Source** | **Dependent Variable** | **Type III Sum of Squares** | **df** | **Mean Square** | **F** | **Sig.** | **Partial Eta Squared** |
| **Corrected Model** | TLR2 | 7.423a | 54 | 0.137 | 2.101 | 0.207 | 0.958 |
| | IFN-γ | 15.380b | 54 | 0.285 | 0.361 | 0.973 | 0.796 |
| | IL-6 | 6.807c | 54 | 0.126 | 1.484 | 0.355 | 0.941 |
| | IL-10 | 6.252d | 54 | 0.116 | 1.574 | 0.326 | 0.944 |
| | TGF-β | 23.284e | 54 | 0.431 | 3.073 | 0.104 | 0.971 |
| | GATA3 | 26.864f | 54 | 0.497 | 2.721 | 0.131 | 0.967 |
| **Intercept** | TLR2 | 212.203 | 1 | 212.203 | 3244.192 | <0.001 | 0.998 |
| | IFN-γ | 185.058 | 1 | 185.058 | 234.772 | <0.001 | 0.979 |
| | IL-6 | 214.378 | 1 | 214.378 | 2523.821 | <0.001 | 0.998 |
| | IL-10 | 222.716 | 1 | 222.716 | 3027.542 | <0.001 | 0.998 |
| | TGF-β | 61.123 | 1 | 61.123 | 435.606 | <0.001 | 0.989 |
| | GATA3 | 53.388 | 1 | 53.388 | 291.964 | <0.001 | 0.983 |
| **gDNA (%)** | TLR2 | 7.423 | 54 | 0.137 | 2.101 | 0.207 | 0.958 |
| | IFN-γ | 15.380 | 54 | 0.285 | 0.361 | 0.973 | 0.796 |
| | IL-6 | 6.807 | 54 | 0.126 | 1.484 | 0.355 | 0.941 |
| | IL-10 | 6.252 | 54 | 0.116 | 1.574 | 0.326 | 0.944 |
| | TGF-β | 23.284 | 54 | 0.431 | 3.073 | 0.104 | 0.971 |
| | GATA3 | 26.864 | 54 | 0.497 | 2.721 | 0.131 | 0.967 |
| **Error** | TLR2 | 0.327 | 5 | 0.065 | | | |
| | IFN-γ | 3.941 | 5 | 0.788 | | | |
| | IL-6 | 0.425 | 5 | 0.085 | | | |
| | IL-10 | 0.368 | 5 | 0.074 | | | |
| | TGF-β | 0.702 | 5 | 0.140 | | | |
| | GATA3 | 0.914 | 5 | 0.183 | | | |
| **Total** | TLR2 | 228.184 | 60 | | | | |
| | IFN-γ | 208.312 | 60 | | | | |
| | IL-6 | 230.167 | 60 | | | | |
| | IL-10 | 238.372 | 60 | | | | |
| | TGF-β | 87.447 | 60 | | | | |
| | GATA3 | 81.889 | 60 | | | | |
| **Corrected Total** | TLR2 | 7.750 | 59 | | | | |
| | IFN-γ | 19.321 | 59 | | | | |
| | IL-6 | 7.232 | 59 | | | | |
| | IL-10 | 6.619 | 59 | | | | |
| | TGF-β | 23.985 | 59 | | | | |
| | GATA3 | 27.779 | 59 | | | | |

a R Squared = 0.958 (Adjusted R Squared = 0.502)

b R Squared = 0.796 (Adjusted R Squared = -1.407)

c R Squared = 0.941 (Adjusted R Squared = 0.307)

d R Squared = 0.944 (Adjusted R Squared = 0.344)

e R Squared = 0.971 (Adjusted R Squared = 0.655)

f R Squared = 0.967 (Adjusted R Squared = 0.612)
